# Tick Innate Immune Responses to Hematophagy and *Ehrlichia* Infection at Single-Cell Resolution

**DOI:** 10.1101/2023.10.02.560565

**Authors:** Abdulsalam Adegoke, Jose M. C. Ribeiro, Ryan C. Smith, Shahid Karim

## Abstract

Ticks rely on robust cellular and humoral immune responses to control microbial infection. However, several aspects of the tick’s innate immune system remain uncharacterized, most notably that of the immune cells (called hemocytes), which are known to play a significant role in cellular and humoral responses toward microbes. Despite the importance of hemocytes in regulating microbial infection, our understanding of their basic biology and molecular mechanisms remains limited. Therefore, we believe that a more detailed understanding of the role of hemocytes in the interactions between ticks and tick-borne microbes is crucial to illuminate their function in vector competence and to help identify novel targets for developing new strategies to block tick-borne pathogen transmission. This study examined hemocytes from the lone star tick (*Amblyomma americanum*) at the transcriptomic level using the 10X genomics single-cell RNA sequencing platform to analyze hemocyte populations from unfed, partially blood-fed, and *Ehrlichia chaffeensis*-infected ticks. Our data exhibit the identification of twelve and nineteen distinct hemocyte populations, respectively, from uninfected and *Ehrlichia*-infected ticks. Our results show a significant increase of clusters representing granulocyte and oenocytoids populations with *Ehrlichia* infection. This work opens a new field of tick innate immunobiology to understand the role of hemocytes, particularly in response to prolonged blood-feeding (hematophagy) and tick-microbial interactions.

**Significance:** The immune response of ticks plays a crucial role in their ability to survive and transmit pathogens. Hemocytes, the primary immune cells in arthropods, are key mediators of tick’s immune defense and provide deep insights into immune responses to microbial infection. However, tick hemocytes’ cellular complexity and heterogeneity have posed challenges for comprehensive characterization. This study employed single-cell RNA sequencing (scRNA-seq) to profile tick hemocytes and elucidate their transcriptional diversity. We identified distinct subpopulations of hemocytes and characterized their unique gene expression profiles. We observed significant variation in immune-related gene expression between hemocyte subpopulations during hematophagy and in response to *Ehrlichia* infection, suggesting specialized functional roles. Our analysis revealed potential marker genes associated with specific hemocyte functions. Our study provides the first comprehensive single-cell atlas of tick hemocytes, shedding light on these cells’ molecular diversity and immune functions. The findings enhance our understanding of tick-host interactions and offer a step toward understanding arthropod immunobiology.

## Introduction

Hematophagous arthropods and insect vectors transmit numerous pathogenic microbes, including bacteria, viruses, and parasites, posing a significant threat to human and animal health worldwide. These blood-feeding ectoparasites have evolved intricate mechanisms to overcome host immune defenses and ensure successful feeding and pathogen transmission. Understanding the cellular components and dynamics of the tick immune system is essential for unraveling the underlying mechanisms of tick-host interactions and developing effective strategies for controlling tick-borne diseases. The immune cells of arthropods or hemocytes serve as the primary line of defense against invading pathogens. In ticks, these versatile immune cells play a vital role in recognizing, encapsulating, and eliminating pathogens acquired during feeding or through vertical transmission (Inoue et al., 2001; Urbanová et al., 2017; Mondekova et al., 2017). Hemocyte mediates several immune processes ranging from phagocytosis, nodulation, melanization, and the release of antimicrobial peptides (Feitosa et al., 2018; Pereira et al., 2001; Fogaça et al., 2006; Kocan et al., 2008; Fiorotti et al., 2022; Adegoke et al., 2023a, Adegoke et al., 2023b)

Tick hemocytes exhibit remarkable heterogeneity and complexity, comprising multiple distinct subpopulations with unique functional attributes. Traditional microscopy approaches have provided valuable insights into the overall composition and morphology of tick hemocytes (Binnington and Obenchain, 1982; Adegoke et al., 2023), leading to the identification of five types: prohemocytes, plasmatocytes, granulocytes, spherulocytes, and oenocytoids (Binnington and Obenchain, 1982; Adegoke et al., 2023). Interestingly, several studies have reported fewer hemocyte types in other tick species (Sonenshine and Roe, 2013; Borovičková and Hypša, 2005). Transcriptional studies of tick hemocytes (Kotsyfakis et al., 2015; Adegoke et al., 2023) have revealed important information regarding the molecular regulation of tick hemocytes during physiological processes such as blood feeding or microbial infection. Nevertheless, these techniques often fail to capture the full spectrum of cellular diversity and functional heterogeneity at the single-cell level, limiting our understanding of the intricate cellular immune responses accompany blood feeding or infection.. Single-cell RNA sequencing (scRNA-seq) has redefined our understanding of hemocyte populations in invertebrate models such as *Drosophila* (Cattenoz et al., 2020; Cho et al., 2020; Tattikota et al., 2020), mosquito (Severo et al., 2018; Raddi et al., 2020; Kwon et al., 2021), silkworm and moth (Feng et al., 2021; Feng et al., 2022), ticks (Rolandelli et al., 2023) and aquatic invertebrates (Koiwai et al., 2021; Pichon et al., 2022; Cui et al., 2022). Applying scRNA-seq in these organisms has led to identifying additional hemocyte complexity and cell types when compared to previous classifications (Raddi et al., 2020; Kwon et al., 2021). In addition, its application has redefined hemocyte complexity in response to pathogen infection (Feng et al., 2022).

This study presents a comprehensive single-cell characterization of the lone-star tick (*Amblyomma americanum*) hemocytes, aiming to unravel the cellular complexity and functional diversity in response to blood feeding and *Ehrlichia* (E) *chaffeensis* infection for the first time. We identified a higher diversity of hemocyte clusters defined by highly expressed marker genes specific to each cluster when hemocytes from the uninfected cohort were compared to *E. chaffeensis*-infected cohorts. The task of correlating the transcriptomic populations from this study with the morphological populations obtained from microscopy and molecular studies in *A. americanum* is still challenging due to the lack of a genome and limited transcriptome and proteome. However, our findings suggest the presence of highly diverse hemocyte populations in *A. americanum*.

## Methods

### Tick rearing and Generation of Ehrlichia-infected ticks

Fully replete *Amblyomma americanum* nymphs were purchased from the Oklahoma State University’s Tick Rearing Facility and maintained at 34℃ and 65% RH under 14:10 h L:D photoperiod till needed. *Ehrlichia* infected adult ticks were generated as previously described (Karim et al., 2012). Briefly, fully engorged nymphs were injected with *E. chaffeensis* (Arkansas strain) using a 32-guage needle fitted to a Hamilton syringe (Hamilton Company, Franklin, MA, USA). All injected ticks were monitored for 25 hours at a temperature of 34℃ to remove dead or non-viable nymphs. Viable nymphs were transferred into an incubator maintained at 34℃ and 65% RH under 14:10 h L:D photoperiod and monitored until they molted into adult male and females. Adult ticks were fed on sheep and removed at the partially-fed phase (50-100mg).

### RNA extraction, cDNA synthesis, and qRT-PCR

Five individual ticks were selected, and tissues dissected as previously described (Karim et al., 2002). RNA was extracted from individual tissue using the Trizol-chloroform separation and isopropanol precipitation method. The quality of the isolated RNA was checked and stored at - 80℃ until use. Complementary DNA (cDNA) was synthesized from the isolated RNA as previously described (Bullard et al., 2016). *Ehrlichia* infection was confirmed from individual tissues by nested PCR amplification of the 16S rRNA 389-bp from the cDNA (Dawson et al. 1994, Steiner et al., 1999). The amplified products were analyzed on a 2% agarose gel stained with SYBR safe and imaged using the Gel-documentation system (Bio-Rad, Hercules, CA).

### Single hemocyte encapsulation

Three independent cohorts of adult female lone star ticks (*Amblyomma americanum)* representing unfed, blood-fed, and *Ehrlichia*-infected (Arkansas strain) ticks were prepared for hemolymph collection. Hemolymph was collected from these three cohorts using our published method with slight modifications (Adegoke et al., 2023). Collected hemolymph was resuspended in Leibovitz’s L-15 Medium supplemented with 50% BSA. Hemocyte suspension was filtered through a 40 µm cell strainer and centrifuged at 500 g for 3 min at 4°C. The hemocyte pellet was washed and centrifuged five times using 1 mL of ice-cold Leibovitz’s L-15 Medium supplemented with 50% FBS. The final pellet was resuspended in 200 µL PBS supplemented with 5% BSA.

### Library preparation and sequencing

Single cell samples were assessed for proper quality control metrics prior to library preparation. All samples were quantitated through the Invitrogen Countess 3 FL Automated Cell Counter (Invitrogen, Thermo Fisher Scientific, Waltham, MA, USA) using ReadyCount Green/Red Viability Stain (ReadyCount Green/Red Viability Stain) per manufactures’ protocol with GFP filter. Samples were all greater than 90% viable with cell density averaging 1000 cells/ul. A total of 10000 cells per sample were used for downstream library preparation. Single cell samples were processed through the 10X Genomics Chromium Single Cell 3’ protocol v3.1 per manufactures’ instruction. Single cell droplets (GEMs) were produced using the 10X Genomics Chromium Controller and barcoded for sample identification. Final libraries were validated on the Qiagen QIAxcel DNA High Sensitivity Bioanalyzer (QIAGEN, Germantown, MD, USA)), to visualize proper insert size, and concentration was calculated using the KAPA Illumina Quantification Kit (Kapa Biosystems, Wilmington, MA, USA) on the Bio-Rad CFX96 thermal cycler (Bio-Rad Inc., Hercules, CA, USA). Pooled libraries were sequenced on the Illumina NextSeq 2000 per 10X Genomics specifications (28×91 cycles).

### Quality control and data processing

The obtained raw single-cell sequencing dataset was analyzed for quality statistics using the official 10X Genomics analysis software Cell Ranger (10x Genomics, Cell Ranger 3.1.0). The forward and reverse reads were uploaded to the cell ranger cloud platform, and reference alignment was carried out on a custom-made reference file. When analyzing this data, no reference genome for A. americanum is available. To navigate this technical hurdle, we created a pseudo-genome using publicly available transcriptomes to generate contigs that were “stitched” with 50 N spacers in a pseudo-chromosome fasta (fa) file from which a general feature format (gtf) file was created (Supplemental File 1). The short reads used for this assembly have the NCBI Short Reads Archive (SRA) accessions SRR1740607, SRR1740608, SRR1740609, SRR1740611, SRR1027751, SRR1027762, SRR1027471, SRR1027473, SRR1027474, SRR1027475, SRR1027476, SRR1027477, SRR1027479, SRR1027481, SRR1027483, SRR1027485, SRR1755762, SRR1756275, SRR1756327, SRR1756363, SRR1756401, SRR1756447, SRR1765066, SRR1755912, SRR1755757, SRR1765232, SRR1755874, SRR1755882, SRR1755921, SRR1755945, SRR1756279, SRR1756256, SRR1756263, SRR1756048, SRR1755913, SRR1756309, SRR1765128, SRR1765136, SRR1765151, SRR1755789, SRR1755723, SRR1765137, SRR1756448, SRR1756420, SRR1756431, SRR1756433, SRR1756139, SRR1755902, SRR1755922, SRR1756449, SRR1765217, SRR1755736, SRR1755893, SRR1765079, SRR1765248, SRR1755894, SRR1756287, SRR1765067, SRR1756059, SRR1755946, SRR1756101, SRR1756251, SRR1756280, SRR1756069, SRR1755923, SRR1764971, SRR1764999, SRR1755753, SRR4416250, SRR4416251, SRR4416252, SRR4416253, SRR4416254, SRR4416255, totaling 7,3 x 10^10^ bases. After trimming bases with low quality (less than 30) or matching Illumina primers, the reads were converted from fastq to fasta and normalized using the program insilico_read_normalization.pl from the Trinity package, allowing maximum coverage of 100 and smaller coverage of 5 (Haas, Papanicolaou et al. 2013). The normalized file served as input to the Abyss assembler (Simpson, Wong et al. 2009) (with -k values of 20 – 90 in 10 x increments) and to the Trinity assembler. The resulting assemblies were combined into a single fasta file and clustered to 98 % identity with the program CD-HIT-ESTs (Li and Godzik 2006). The BUSCO version 5.0 (Simão, Waterhouse et al. 2015) program running the Arachnida dataset against the assembled transcriptome file indicated 89.0% complete BUSCOs, of which 76.0% were complete and single copy, 13.0% were duplicated and 2.1% were fragmented, while 8.9% were missing out of 2,934 BUSCO genes. Since the assembled transcripts included hemocyte libraries, we estimate our gene coverage for the hemocyte-expressed transcripts was better than 90%. Finally, the CDSs were mapped to a hyperlinked Excel spreadsheet (Supplemental File 2) containing matches to several databases, allowing an estimate of their functional classification.

### Dimensionality reduction and cell clustering

The output matrices generated from the Cell Ranger software were loaded into the Seurat R package for cell clustering and dimensionality reduction analysis (Hao et al., 2021). Data were normalized using the ‘LogNormalize’ function, and the variations between cells were regressed by counting the number of target molecules in each cell, Unique Molecular Identifiers (UMI). Next, the scaled data underwent Principal Component Analysis (PCA) to reduce dimensionality. The JackStrawPlot function was utilized to examine the distribution of P-values. Among the PCA results, the principal component (PC) with the most significant statistical significance (p <10-5) was chosen for subsequent clustering and cluster analysis (Chung and Storey, 2015). Leveraging the identified PCs, cells were clustered and organized using the ’FindClusters’ function. To further investigate data structures and cell trajectories, t-SNE and UMAP were employed, enabling separate visualization and exploration (McDavid et al., 2013; Perrin and Zuccon, 2018).

### Marker gene identification

The Seurat functions ’FindAllMarkers’ and ’roc’ were employed to identify marker genes associated with the obtained cell clusters. The marker gene refers to a set of genes used to identify and characterize identified hemocyte clusters within the cohorts. This analysis involved testing the cluster-enriched genes for significance. Subsequently, cluster-specific marker genes were subjected to additional screening criteria: they had to be expressed in over 50% of cells within a given cluster and exhibit an average natural log-fold change greater than 0.5. Multiple replicate analyses were conducted using either the same or different parameters to ensure the robustness and accuracy of the marker genes.

### Pseudotemporal ordering of cells

Slingshot was used to comprehensively examine cell differentiation and fate. The data was transferred to the slingshot package by leveraging information from the identified cell clusters and differentially expressed genes. Subsequently, genes associated with cell differentiation and cell fate were selected to define the progression of cellular differentiation. Slingshot effectively reduced the dimensionality of the data to two dimensions and arranged all cells in a meaningful order. Genes exhibiting similar expression patterns, potentially indicating shared biological functions, were grouped together. The ’learning-graph’ function was employed to calculate cell trajectories, providing insights into the progression of cellular differentiation.

## Code Availability

The generated data were analyzed using the 10X genomics Cell Ranger v7.1 (10x Genomics, Cell Ranger 3.1.0). Data analysis and visualization were carried out following the Seurat – Guided Clustering Tutorial (https://satijalab.org/seurat/articles/pbmc3k_tutorial.html)in R studio using all default settings.

## Data Availability

The raw datasets generated in this study are deposited in the NCBI Transcriptome Shotgun Annotation (TSA) repository under the accession number ########. The pseudo-genome and pseudo-GTF files (Supplemental file 1) can be downloaded from https://proj-bip-prod-publicread.s3.amazonaws.com/transcriptome/Amb_americanum_pseudogenome/amb_amer_pseudo_fa_gtf.zip Supplemental file 2, containing the assembled and annotated CDS can be downloaded from https://proj-bip-prod-publicread.s3.amazonaws.com/transcriptome/Amb_americanum_pseudogenome/Amb_amer-s2.zip

## Results

## Single Cell RNA-seq of hemocytes from A. americanum

To examine hemocyte heterogeneity by scRNA-seq, isolated hemocytes from unfed, partially fed, and partially fed-*Ehrlichia chaffeensis* infected *A. americanum* ticks were subjected to the 10X Chromium Single Cell 3’ sequencing platform. Approximately 27,130 cells (unfed: 6445; partially fed: 5847; partially fed *Ehrlichia*-infected: 14838) were captured using the 10X Genomics Chromium microfluidic technique and submitted to RNA sequencing. After standard pre-processing, a total of 26,040 cells covering 5870 cells from unfed, 5427 cells from partially blood-fed, and 14,744 cells from *Ehrlichia*-infected ticks were detected. A sequencing depth of ∼14,000 reads per cell at a median sequence saturation of 69.6% (unfed), 67.9% (blood-fed), and 66.5% (*Ehrlichia* infected) across libraries were achieved. We obtained a median Unique Molecular Identifiers (UMI) and gene counts of 835 and 3181 from uninfected cohorts, 915 and 3478 from partially fed cohorts, and 582 and 2281 from partially fed-*Ehrlichia*-infected cohort (Fig. 1A-C), respectively. Principal component linear dimensional reduction and the Uniform Manifold Approximation and Projection (UMAP) tool were used to visualize the filtered datasets (Becht et al., 2019; Stuart et al., 2019). Prior to unsupervised clustering, we applied the Seurat Integration batch correction method on all three cohorts. Seurat’s unsupervised clustering analysis on the transformed cohort grouped hemocytes into fourteen clusters (Fig. 2A) based on similar gene expression profiles (Niu et al., 2020; Kwon et al., 2021; Raddi et al., 2020). For the comparison between the three conditions, each cluster had the following number of cells: Cluster 0 with 3903 cells (14.4%), Cluster 1 with 3727 cells (13.7%), Cluster 2 with 3210 cells (11.8%), Cluster 3 with 3045 cells (11.2%), Cluster 4 with 2846 cells (10.5%), Cluster 5 with 2422 cells (8.9%), Cluster 6 with 1715 cells (6.3%), Cluster 7 with 1452 cells (5.4%), Cluster 8 with 1259 cells (4.6%), Cluster 9 with 1226 cells (4.5%), Cluster 10 with 903 cells (3.3%), Cluster 11 with 799 cells (2.9%), Cluster 12 with 412 cells (1.5%), and Cluster 13 with 207 cells (0.8%) (Fig. 2B). All cohorts appears to have equal proportion of cells represented across clusters, however, cells from *Ehrlichia*-infected cohorts represents more than 50% of cells in clusters 0, 2, 4, 5, 6, 7, 9, 11 and 12 (Fig. 2C and D).

**Figure 1.**
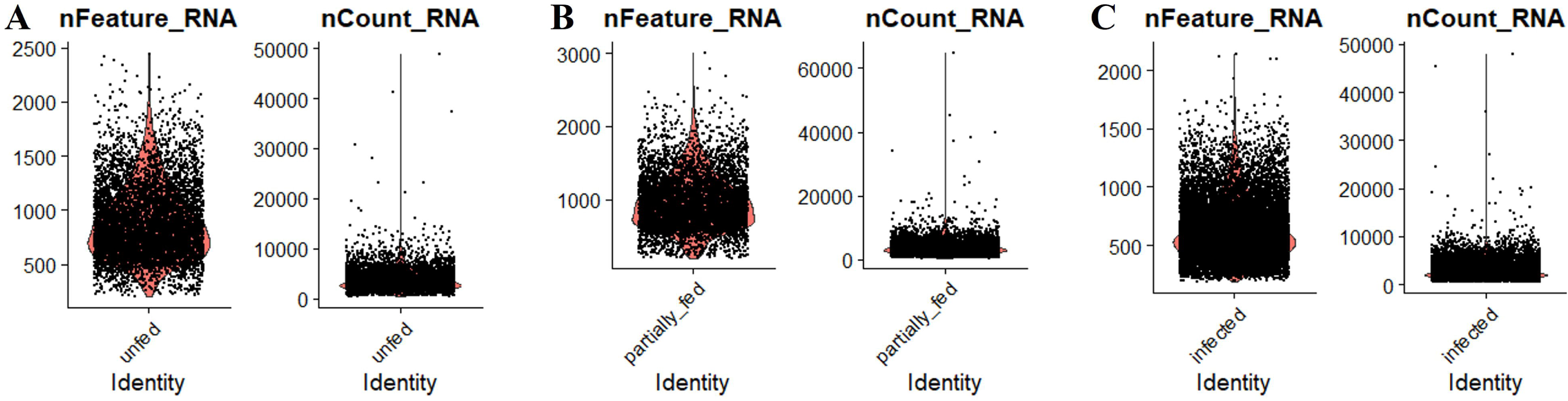
The distribution of the number of transcripts (nFeature_RNA) and genes (nCount_RNA) detected per hemocyte in A) unfed, B) partially fed, and C) *Ehrlichia chaffeensis*-infected cohort.

**Figure 2.**
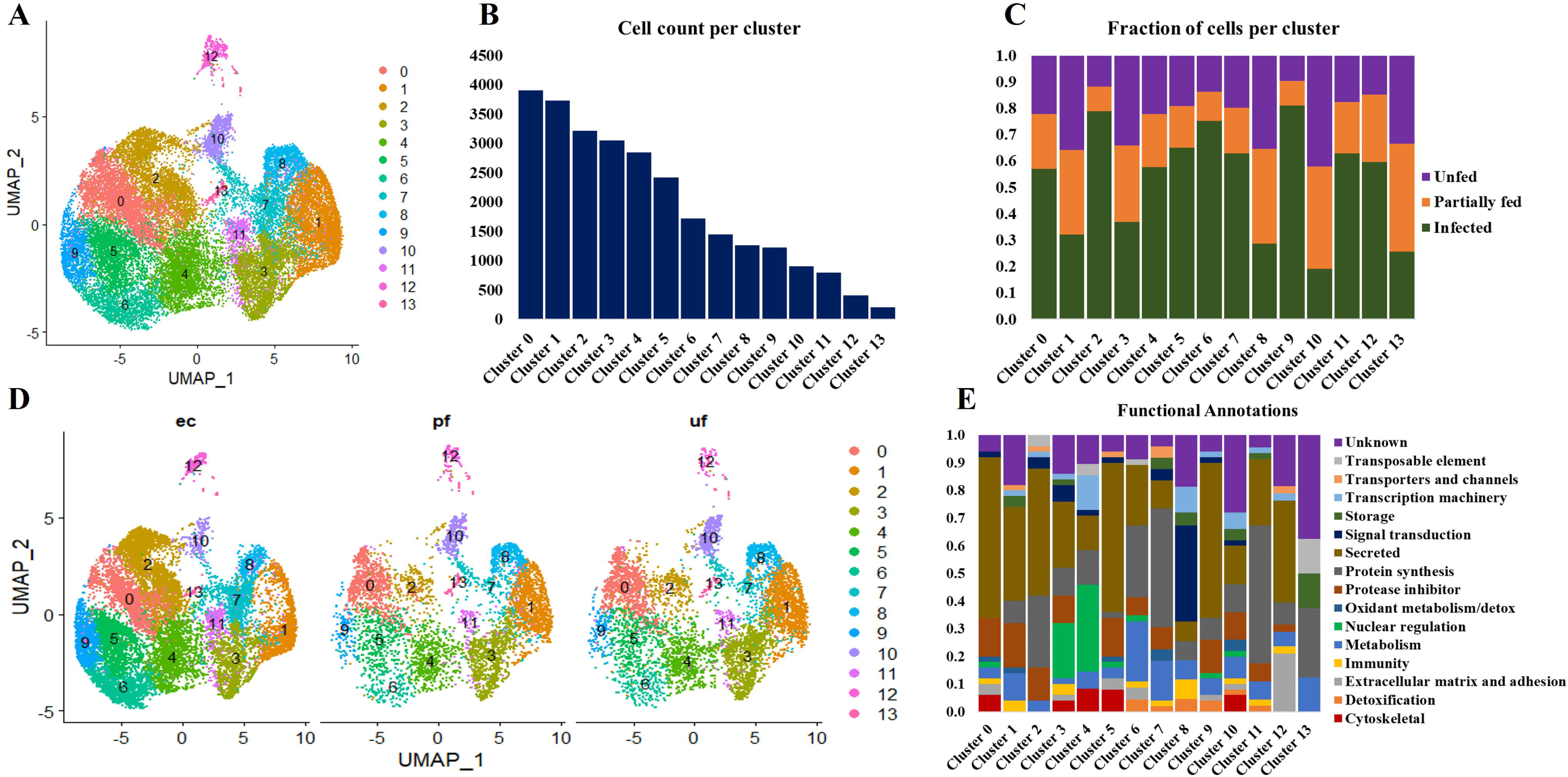
Uniform Manifold Approximation and Projection (UMAP) projection of hemocyte clusters identified from unfed, partially fed and *Ehrlichia chaffeensis* infected *A. americanum* ticks A). Total number of cells in each Seurat cluster B) and fraction of cells represented in each cluster from unfed, partially fed and *Ehrlichia chaffeensis* infected ticks C). UMAP projection of the differences in cluster density in unfed, partially fed and *Ehrlichia chaffeensis* infected ticks and functional annotation of hemocyte clusters using the top 100 DEGs. Dot plot profiling of differentially expressed genes F). The color gradient of dots represents the expression level, while the size represents the percentage of cells expressing any genes per cluster.

### Functional classification of hemocyte clusters

Seurat FindMarker function was used to identify the top 100 Differentially Expressed Genes (DEGs) in each cluster and functional assignment was carried out based on homology search in other tick species, UNIPROT, KOG, PFAM and SMART databases. The identified hemocyte clusters were characterized based on these functional assignments (Fig. 2E, Supplemental File 3).

Clusters 0, 5, and 9 display high expression of secreted genes with roles as protease inhibitors and cytoskeletal functions (Fig. 2E). The marker genes in these clusters include a translation initiation factor 3 (14835), a metalloprotease (116223), lipocalins (16064, 146185, 90157), hemocytin (90421) and a serine protease inhibitor (137223), which are all involved in cellular and humoral immune response (Fig. 2F).

The cells in cluster 1 express secreted genes involved in protein synthesis, protease inhibitors, metabolism, and immune response (Fig. 2E). The identified markers in this cluster include Inositol 1,4,5-trisphosphate receptor (120699; 120701), adenylyl cyclase (107091), serine protease inhibitor (31360), chymotrypsin inhibitor (105147), and carboxypeptidase inhibitor (766650). Inositol is involved in phagocytosis, and its expression induced via serotonin modulation. At the same time, serpin, chymotrypsin, and carboxypeptidase inhibitors have diverse roles in humoral immunity and blood digestion (Fig. 2F).

Cluster 2 shares a similar functional expression of genes as in cluster 1 (Fig. 2E); however, it is marked with high expression of marker genes such as mucin (459841) and cystatin (9505, 22377), which are all secreted proteins regulating immune response and chitinase (187500), which is involved in wound healing (Fig. 2F).

Cluster 3 and 4 are enriched with genes with functions in nuclear regulation and cytoskeletal functions such as several histones (129995, 9657, 153147, 188513), GTPase-activating protein (81855, 77010), laminin (70101) and kinesin-like protein (195826) (Fig. 2E). These genes are involved in the regulation of cell cycle, gene expression, cytoskeletal organization, and signal transduction. They are also components of the extracellular matrix (ECM) and basement membranes (BMs) and perform roles ranging from cell adhesion to cell-to-cell migration (Fig. 2F).

Cluster 6 is enriched with secreted genes related to protein synthesis, protease inhibitor, metabolism, and detoxification (Fig. 2E). The marker genes in this cluster include a glutathione S-transferase (175001), mitochondrial NADH: ubiquinone oxidoreductase (82400), heat shock protein (58400). These genes are involved in the production of ROS and the regulation of oxidative stress. ROS is an essential signaling molecule in wound healing cell differentiation and innate immunity (Fig. 2F).

Cluster 7 showed high expression of genes related to protein synthesis, protease inhibitor, and metabolism (Fig. 2E). The highly expressed marker genes include cytochrome_P450 (1455319), low-density lipoprotein receptor (58886) and cystatin-B (145319). These genes are involved in the cellular stress response, activation of lysosomal enzymes, lipoprotein metabolism, and protease degradation (Fig. 2F).

Cluster 8 showed high expression of genes related to signal transduction, transcription machinery, immunity, and protein synthesis (Fig. 2E). Such genes identified in this cluster include intersectin (102281), translation initiation factor IF-2 (1495781), hemocytin-like (113156) and calpain-b (98929). These genes are important regulators of cell transport, apoptosis regulation, and microbial binding and aggregation (Fig. 2F).

Cluster 10 is enriched for secreted genes and genes cytoskeletal, transcriptional machinery, protein synthesis, protease inhibitor, and metabolic functions (Fig. 2E). The marker genes in this cluster include cathepsin-L (89264), transforming growth factor-beta-induced protein ig-h3 (51103), soma ferritin (154941), a putative peptidase inhibitor (1018222), and fibronectin (106951). These marker genes are involved in protein degradation, cell differentiation, transport, and cellular differentiation (Fig. 2F).

Cluster 11 is enriched with genes involved in protein synthesis and secretory function, such as keratinocyte-associated protein 2 (125644), carboxypeptidase inhibitor (76650), and serine protease inhibitor (31360) (Fig. 2E). The presence of these genes will suggest the role of this cluster in cytoskeletal binding, encapsulation, and tissue remodeling during repair (Fig. 2F).

Cluster 12 has high expression of secreted genes and genes involved in protein synthesis and extracellular matrix and adhesion (Fig. 2E). The marker genes in this cluster include cuticle protein (42855), transglutaminase substrate-like (181535), a glycine-rich protein (134628, hemicentin (933328), and reeler domain-containing protein (28024). The functional role of these genes involves the formation of the extracellular matrix and cuticle development, cell differentiation, and the formation of the peritrophic matrix (Fig. 2F).

Cluster 13 has the least expressed number of genes and is not of hemocyte origin. However, functional enrichment showed the few genes detected are involved in storage, protein synthesis, and metabolic processes (Fig. 2E and F).

### Cell lineage analysis

We used slingshot to perform pseudotime analysis from the obtained hemocyte clusters to investigate the potential differentiation trajectory and reveal lineage relationships between the identified clusters. Seven hemocyte lineages were predicted. The first lineage starts from cluster 0 through clusters 5, 6, 4, 11, 7 and ends at cluster 1. The second lineage originates from cluster 0 through clusters 5, 6, 4, 11, 13 and ends at cluster 10. Lineage 3 originates from cluster 0 through clusters 5, 6, 4, 11, 13 and ends at cluster 12. Lineage 4 originates from cluster 0 through clusters 5, 6, 4, 11, 7 and ends at cluster 8. Lineage 5 originates from cluster 0 through clusters 5, 6, 4, 11 and ends at cluster 3. Lineage 6 originates from cluster 0 through clusters 5 and ends at cluster 9. Lineage 7 runs from cluster 0 and ends at cluster 2 (Fig. 3).

**Figure 3.**
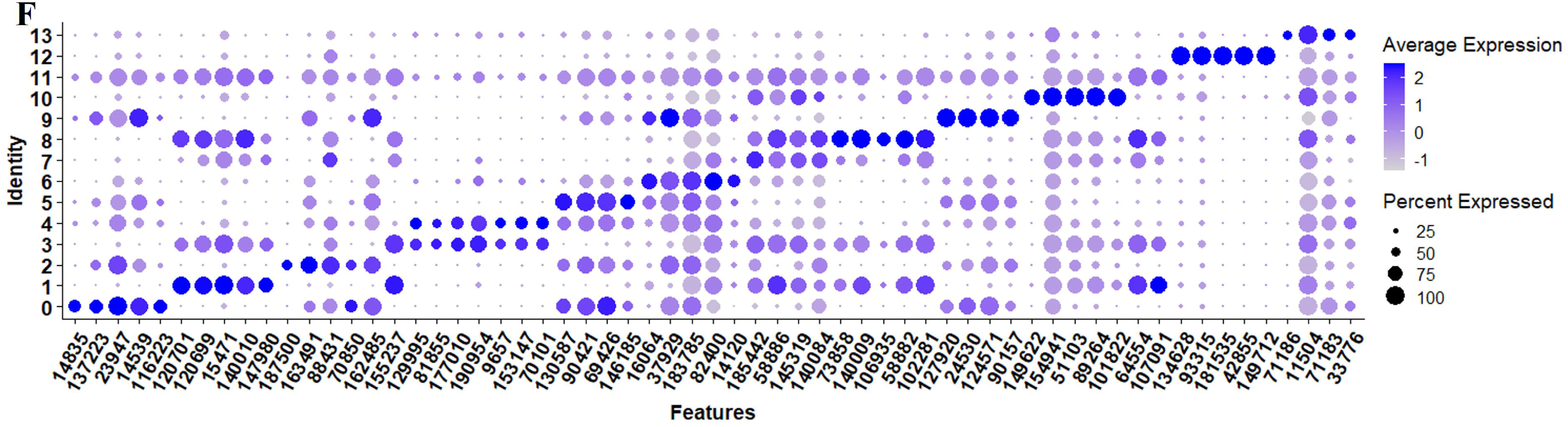
Lineage analysis using SlingShot. In total, seven lineages were identified which defines the trajectory of each the defined clusters. Lineage 1 (L1) includes clusters 0. 5, 6, 4, 11, 7 and 1. Lineage 2 (L2) includes clusters 0, 5, 6, 4, 11, 13 and 10. Lineage 3 (L3) includes clusters 0, 5, 6, 4, 11, 13 and 12. Lineage 4 (L4) includes clusters 0, 5, 6, 4, 11, 7 and 8. Lineage 5 (L5) includes clusters 5, 6, 4, 11 and 3. Lineage 6 (L6) includes clusters 0, 5 and 9. Lineage 7 (L7) includes clusters 0 and 2.

Considering the lack of known hemocyte markers, we calculated and searched for highly expreesed genes over pseudotime. However, we did not observe a unique association between the identified genes and distinct clusters or lineages, rather a group with multiple clusters (Fig. 4A-D). We identified a putative soma ferritin gene (154941) which was differentiated in cluster 10 as part of lineage 2 (Fig. 4A). A putative histone H4 (129995) is enriched in clusters 3, 4 and 11, which are components of lineage 5 (Fig. 4B). The expression of a lipocalin-5 (16064) in clusters 5, 6, 9 is high compared to clusters 1, 3, and 11 (Fig. 4C). A laminin (70101) transcript is enriched in clusters 3 and 4 (Fig. 4D). In addition, differentially expressed genes that changes across lineages were identified. The genes include a ribosomal protein S16 (88431), a hypothetical secreted protein (23947), a translation initiation factor-2 (14835), a transposon (188473), lipocalin-5 (16064) and a chitinase-like lectin (187500) (Fig. 5A-F). Seven lineages with shared clusters amongst multiple lineages indicate that many hemocyte populations display pluripotency and can differentiate into different cell types.

**Figure 4.**
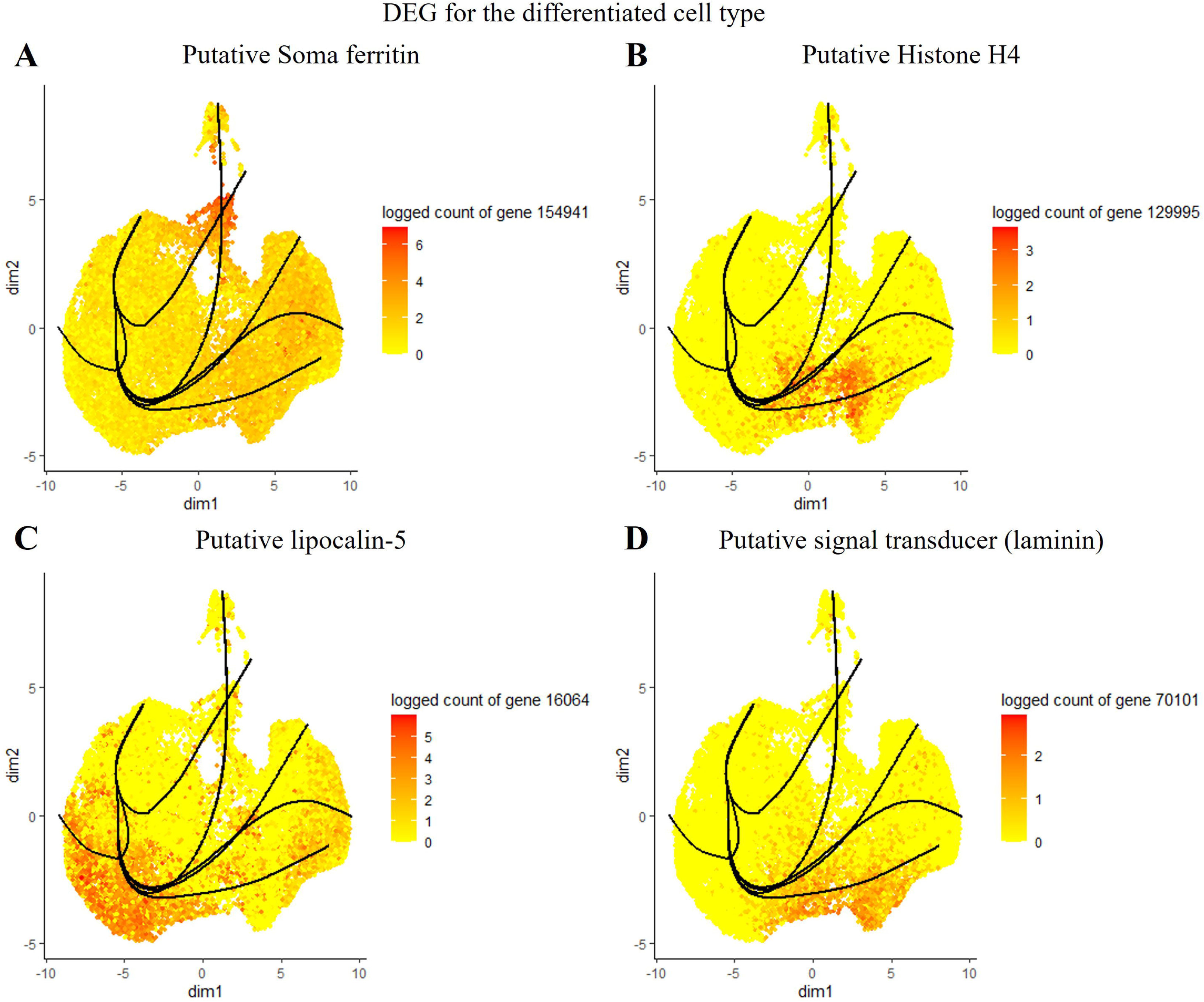
Expression of top genes including putative soma ferritin A), putative histone h4 B), lipocalin-5 C) and laminin D) as their expression changes across pseudotime.

**Figure 5.**
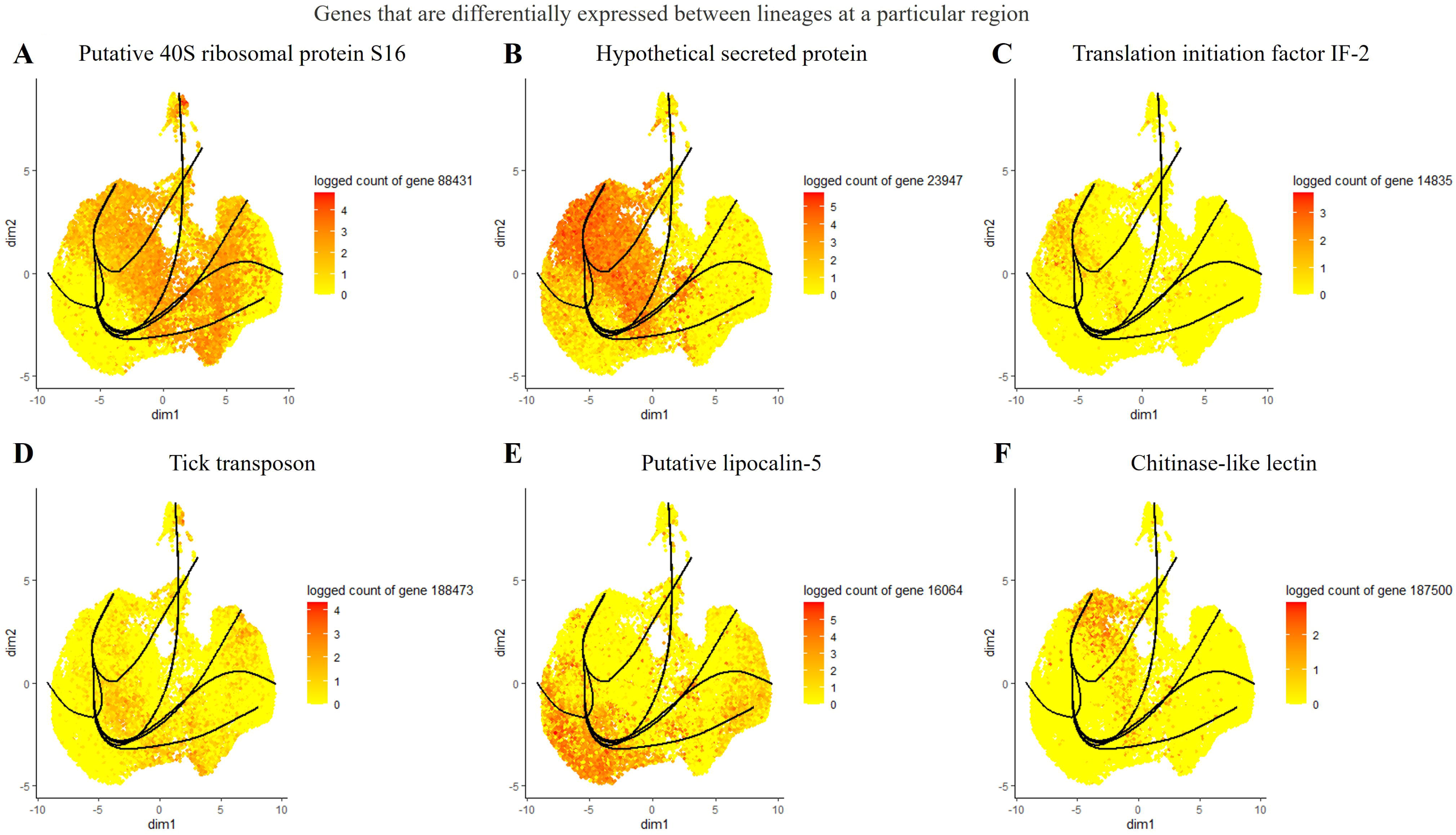
List of differentially expressed genes between lineages at different branches. The top 6 genes include a putative 40S ribosomal protein S16 A), a hypothetical protein B), translation initiation factor IF-2 C), tick transposon D), lipocalin-5 E), and chitinase-like-lectin

## Discussion

The innate immune system of hematophagous arthropods is essential to their response to commensal and pathogen infection. We recently characterized the morphological changes and molecular responses of tick *A. maculatum* and *A. americanum* hemocytes to *R. parkeri* and *E. chaffeensis* infection (Adegoke et al., 2023a, 2023b). However, technical challenges and the paucity of reagents related to characterized molecular markers limit functional studies on these cell types. In the current study, we sequenced hemocytes isolated from *A. americanum* ticks under unfed, partially fed, and *E. chaffeensis*-infected conditions using a single-cell approach. Our result showed extensive heterogeneity in hemocyte populations, with significant changes observed in hemocytes infected with intracellular tick-borne pathogens, *E. chaffeensis*.

This study identified 14 distinct hemocyte clusters, and the most notable changes in hemocyte population and cluster proportion were seen with *E. chaffeensis* infection. The clusters with a fraction of cells induced by *E. chaffeensis* infection were notable for their expression of genes involved in cytoskeletal functions, nuclear regulation, protein synthesis, metabolism, detoxification, and extracellular matrix secretion. Interestingly, these clusters are shared across three different lineages, 4, 6, and 7, suggesting a common ancestry among the cells from these clusters. The genes expressed in these clusters are involved in microbial recognition and agglutination, cellular differentiation and proliferation, and response to oxidative stress.

Considering the difficulty in functionally characterizing the role of the identified genes in hemocyte functions, we focused on those genes that are highly abundant in clusters enriched in *E. chaffeensis-*infected cohorts. These genes include hemocytin, chitinase-like proteins, serine-protease inhibitors, lipocalins, cystatins, heat shock proteins, and glutathione s-transferase. In addition, we identified genes that are involved in regulating cellular immune response and cellular differentiation. These include histones, laminins, GTPase, and kinesin.

Hemocytin, an ortholog of *Drosophila* hemolectin and human von Willebrand factor, is an adhesive protein released from the granules of granulocytic hemocytes via exocytosis and is the first step in the formation of nodules after infection (Arai et al., 2013; Ratcliffe and Gagen, 1977; Otuka et al., 2023). Chitinase-like proteins (CLPs) belong to the glycoside hydrolase family 18 (GH18). CLPs expression upregulated in *Drosophila* following injury, parasitic infection and bacterial infection by inducing clot formation and wound healing (Irving et al., 2001; De gregorio et al., 2001; Kucerova et al., 2015; De gregorio et al., 2002). Serine-protease inhibitors serve as pattern recognition proteins and are also components of the prophenoloxidase (proPO)-activating system, a major humoral defense mechanism in arthropods (Jiravanichpaisal et al., 2006; Iwanga and Lee, 2005; Cerenius and Söderhäll, 2004; Cerenius et al., 2008). The role of these genes as components of the immune response and their expression in hemocyte clustersenriched in *E. chaffeensis* infected cohorts is indicative of their contribution to the immune response during *E. chaffeensis* infection. Considering the role of these genes in microbial binding, the clusters in which they are enriched (0, 5, and 9) could be regarded as involved in pattern recognition.

Lipocalins are also involved in the transport of small molecules across biological barriers. No direct evidence exists for their role in hemocyte-mediated immunity or their hemocyte-specific expression; however, an *Anopheles (A) gambiae* lipocalin serves as a carrier molecule for a hemocyte differentiator factor, which subsequently leads to immune priming (Ramirez et al., 2015). Alignment of the lipocalin sequences revealed that 67% to 99% amino acid identify with other tick lipocalins (Supplemental File 2, Fig. 1 and Sheet 4) containing the histamine binding motif found in tick salivary proteins. The top hits belonged to sequences from *A. americanum, A. sculptum, A. aureolatum, R. haemaphysaloides,* and *A. maculatum* (Supplemental File 3, Fig. 1 and Sheet 4). It is interesting to note that the three *A. maculatum* (Amac-hemSigP-446830, Amac-hemSigP-446831, and AmHem-446832) transcripts were hemocyte-specific transcripts from *A. maculatum* (Adegoke et al., 2023a). Tick lipocalins perform immunomodulatory roles during tick feeding by scavenging and binding histamines and other biogenic amines (Paesen et al., 2000; Mans and Ribeiro, 2008). Tick lipocalins are functionally diverse, with little sequence identified across tick species. This diversity is also specie dependent as certain tick species, such as *A. maculatum, A. americanum,* reportedly have 170 and 200 lipocalin coding sequences, while species like *H. longicornis* and *I. persulcatus* have just 1 and 12 lipocalin CDS in their genomes (Ribeiro et al., 2023). In *A. americanum,* 200 lipocalin coding sequences are present, constituting approximately 27% of the whole transcriptome in the salivary gland (Karim and Ribeiro, 2015). Similarly, the expression of multiple lipocalin transcripts changes throughout tick feeding, suggesting a time-dependent specificity even in the same tick species (Bullard et al., 2016). Lipocalin expression also changes in response to pathogen infection (Ribeiro et al., 2005). A lipocalin isolated from *Ornithodoros savignyi* showed distinct expression in the hemocytes and midgut following bacterial infection and blood-feeding, respectively, suggesting a tissue-dependent role for lipocalins (Cheng et al., 2010). Tick lipocalins reportedly bind to serotonin and serotonin-binding proteins (Mans et al., 2008; Mans and Ribeiro, 2008; Sangamnatdej et al., 2002). Serotonin and serotonin receptors are responsible for the phagocytic response seen in *Drosophila* and the caterpillar, *Pieris (P) rapae* (Qi et al., 2017) and also in the aggregation of hemocytes and nodule formation in *B. mori* (Otuka and Sato, 2023). The affinity of tick lipocalin towards serotonin or other serotonin-binding proteins would suggest their role in defense response against pathogens with their high expression of multiple lipocalin transcripts in clusters enriched in *E. chaffeensis* infected cohorts.

Cystatins are stored inside the granules of granulocytic hemocytes and possess strong AMP activities against Gram-negative bacteria (Agarwala et al., 1996). The cystatin transcripts from this study were 33-71% identical with previously studied cystatins from other tick species (Suplemental file 2, Fig. 2 and Sheet 5), with the most identity has been salivary cystatin-L from *Hyalomma asiaticum*, *R. sanguineus, R. microplus, Dermacentor (D) silvarum,* and *D. andersoni* (Suplemental file 2, Fig. 2 and Sheet 5). Cystatins are a family of proteins that play important roles in regulating protease activity by inhibiting cysteine proteases, such as Cathepsins, in various biological processes. In ticks, cystatins have unique properties and functions primarily related to tick feeding and evasion of the host immune response. Tick cystatins are essential for the tick’s ability to feed on the host’s blood and inhibit proteases involved in host immune responses (Karim et al., 2005; Kotsyfakis et al., 2007; Parizi et al., 2020; Lu et al., 2020; Kotál et al., 2021). Similarly, ticks secrete cystatins during tick-borne pathogen infection and colonization. Tick cystatins can inhibit cysteine proteases produced by pathogens, as demonstrated in *R. microplus* during *B. microti* colonization (Wei et al., 2020). The expression of multiple cystatin transcripts was upregulated in *R. microplus* salivary gland infected with *Theileria equi* (Paulino et al., 2021). In addition to their presence in salivary gland and midgut, tick hemocytes express high levels of cystatins, suggesting a role in innate immune response (Lu et al., 2014; Kotsyfakis et al., 2015; Zhou et al., 2006). The remarkable diversity and tissue specificity of tick cystatins contribute to their multifaceted roles in tick physiology. Negative regulation of the tick immune system has been described for tick cystatins. Knockdown of *Rmcystatin,* a cystatin from *Rhipicephalus (R) microplus,* rendered ticks immunocompetent upon bacterial infection (Lu et al., 2014). A similar observation was made in the pacific oyster, *C. gigas,* where an enhanced bacterial clearance was detected following the knockdown of the cystatin *CgCytA* (Mao et al., 2018). Our data identified cystatin as a marker in two clusters enriched in *E. chaffeensis-*infected cohorts. *E. chaffeensis* may induce cystatin expression to evade immune response since cystatin can inhibit several proteases found in hemocyte granules (Haves-Zburof et al., 2011).

The expression of laminins, histones, kinesins, and GTPase is unique to clusters 3 and 4. However, while all three cohorts share relatively equal proportions of cells in cluster 3, *E. chaffeensis* infected cohorts have a higher fraction of cells from cluster 4 than the unfed and partially fed cohorts. Histones are indispensable for the organization and architecture of chromatin, and their post-translational modifications play a vital role in gene regulation (Parseghian and Luhrs, 2006). However, they also possess antimicrobial activities against bacteria, fungi, viruses, and protozoa (Kawasaki and Iwamuro, 2008). In vertebrates, they act as damage-associated molecular patterns (DAMPs) and bind to pattern recognition receptors (PRRs), which eventually trigger the innate immune response (Li et al., 2022). Histones are components of extracellular traps (ETs) granulocytes produced upon microbial infection (Nascimento et al., 2018). Granulocyte-produced extracellular traps of *Periplaneta (P) americana* comprised DNA and histones, and the ETs contribute to granulocyte-mediated nodulation following bacterial infection (Nascimento et al., 2018). Laminins are components of extracellular matrix (ECM) and basement membranes (BMs). They interact with cell-surface molecules and other ECM components and perform roles ranging from cell adhesion to cell-to-cell migration. In *Drosophila*, laminins are produced by embryonic hemocytes, which are required for hemocyte migration (Sánchez-Sánchez et al., 2017). Laminins also play vital roles in hemocyte homeostasis, as silencing a laminin receptor decreased the hemocyte population in *Penaeus (Litopenaeus) vannamei* (Charoensapsri et al., 2015). GTPase plays a crucial role in biological processes such as cell cycle regulation, gene expression, cytoskeletal organization, and signal transduction (Hall, 2012). In invertebrates, Rac1, a member of the Rho GTPase family, was highly expressed in the hemocytes of the Chinese shrimp *Fenneropenaeus chinensis,* and the expression was upregulated following viral infection (Chi et al., 2013). Several kinesin-like proteins play a role in chromosomal segregation and congression during meiosis (Zhang et al., 2015; Mayr et al., 2007). Due to their involvement in microtubule dynamics, they are required for efficient phagocytosis (Silver and Harrison, 2011). The high expression profiles of these genes enriched in all cohorts and those enriched in *E. chaffeensis* cohorts, as seen in cluster 3 and cluster 4, respectively, imply that these genes contribute to hemocyte homeostasis and proliferation, as seen with histones and laminins. In addition, they also have additional functions in response to infection and act as migration factors for hemocytes via cytoskeletal reorganization and antimicrobial compounds. When considered together, clusters 3 and 4 may be similar hemocyte populations demonstrating plasticity upon infection.

Glutathione S-transferase and heat shock proteins are highly expressed in cluster 6, enriched in *E. chaffeensis-*infected cohort. Glutathione S-transferases (GSTs) are enzymes that regulate redox homeostasis, detoxification, and innate immune responses against bacterial infection. Multiple sequence alignment of the GST sequence from this study shows identity ranging from 53%-86% identity with most similar sequences coming from *R. microplus, D. andersoni, D. silvarum, D. asiaticum, R. sanguineus, H. longicornis* and *I. scapularis* (Suplemental file 2, Fig. 3 and Sheet 6). GST expression is associated with oxidative stress response and immune response activation (Myers et al., 2018; Ramond et al., 2020). Increased expression of GSTs in ticks is associated with the detoxification of acaricides (Hernandez et al., 2018). Increased hemocyte expression is associated with induction with bacterial components such as LPS or peptidoglycan (Yang et al., 2012) or virus infection (Duan et al., 2013). GTSs also encode enzymes that detoxify Drosophila’s hydrogen peroxide and lipid peroxidases (Singh et al., 2001; Saisawang et al., 2012). ROS production in the hemocytes is required during the early phase of infection for hemocyte activation and bacterial clearance (Myers et al., 2018). A *Drosophila* GST-S1 was reported to induce an increase in larval hemocytes (Stofanko et al., 2008), and the expression of GST-S1 was also increased in hemocytes during early metamorphosis (Regan et al., 2013), suggesting a functional role in regulating hemocyte proliferation. The expression of tick GSTs is associated with blood feeding in response to elevated oxidative stress. Breakdown of blood meals results in high levels of ROS, thus acting as a signal transducer for the expression of GSTs (Daniel, 1993; Dreher-Lesnick et al., 2006). Increased expression of tick GSTs during blood meal is shown in *I. ricinus* and *R. microplus* (Rosa de Lima et al., 2002; Rudenko et al., 2005). In the midgut, transport, and intracellular digestion of heme in the midgut epithelial results in continuous ROS production throughout feeding, resulting in the upregulation of multiple GST transcripts (Perner et al., 2016). The expression of GST in tick salivary gland is seen towards the late stage of feeding, which coincides with salivary gland degeneration and massive release of intracellular salivary proteins into the salivary ducts (Tirloni et al., 2015; Radulović et al., 2014). Other studies have also reported multiple GST transcripts in the salivary gland, ovaries, and fat bodies; however, these GST transcripts gradually reduce as ticks move towards full engorgement (Hernandez et al., 2018). Heat shock proteins (HSP) are associated with elevated stress levels and act as chaperones during high protein turnover or degradation. In invertebrates, elevated levels of HSPs in hemocytes are associated with increased oxidative stress, usually associated with wounding or infection (Chakrabarti et al., 2020; Wrońska and Boguś, 2020). Elevated levels of reactive oxygen species (ROS) and associated oxidative stress during melanization or clotting response to sterile or contaminated wounds (Krautz and Arefin, 2014). The role of both GST and HSP in stress metabolism and cell protection during both blood feeding and infection makes them an integral component of the immune response, further emphasizing their high expression in *E. chaffeensis-infected* cohorts.

In conclusion, we used single-cell transcriptome to delineate hemocyte heterogeneity in a blood-feeding arthropod. We identified 14 unique hemocyte clusters and revealed changes in cluster population induced by *E. chaffeensis*. The findings here serve as a significant resource to generate new knowledge to understand tick immune biology and their responses to microbial infection.

### Limitations

This study has led to the identification of several hemocyte-derived genes that can be used to define hemocyte populations. Using pseudotime and trajectory analysis has identified several potential genes that will be useful in determining hemocyte pluripotency into other hemocyte types. However, the lack of an annotated *A. americanum* genome and hemocyte transcriptome limited our ability to make conclusive inferences.

## Data Availability Statement

The datasets presented in this study are deposited in the Sequence Read Archive (SRA) repository under the accession number.

## Ethics Statement

The animal study was reviewed and approved by Protocols for tick blood-feeding were approved by the University of Southern Mississippi’s Institutional Animal Care and Use Committee (USMIACUC protocols #15101501.3 and 17101206.2).

## Acknowledgments

This work utilized the computational resources of the NIH HPC Biowulf cluster (http://hpc.nih.gov).

## Author contributions

Conceptualization: AA, RCS, SK; Data Curation: AA, JMR, SK; Formal analysis: AA, JMCR, RCS, SK; Funding acquisition: SK; Investigation: AA, JMCR, RCS, SK; Methodology: AA, JMCR, RCS, SK; Project Administration: SK; Resources: JMCR, RCS, SK; Supervision: SK; Writing original draft: AA, SK; Writing, reviewing & editing: AA, JMCR, RCS, SK

## Supplementary Figures

Figure 1: The neighbor-joining phylogenetic tree from the aligned lipocalin sequences following 1000 bootstraps. The bar at the bottom represents 20% amino acid diversity. The numbers at the nodes indicate the percentage bootstrap support.

Figure 2: The neighbor-joining phylogenetic tree from the aligned cystatin sequences following 1000 bootstraps. The bar at the bottom represents 20% amino acid diversity. The numbers at the nodes indicate the percentage bootstrap support.

Figure 3: The neighbor-joining phylogenetic tree from the aligned glutathione s-transferase sequences following 1000 bootstraps. The bar at the bottom represents 20% amino acid diversity. The numbers at the nodes indicate the percentage bootstrap support.

## Supplementary Tables

Sheet 1: NCBI short reads from short reads archive (SRA) and their accessions used for transcripts assembly

Sheet 2: Base statistics of the short reads used

Sheet 3: Combined coding sequences matched to publicly available databases

Sheet 4: Multiple alignment of lipocalin protein sequences using Constraint-based Multiple Alignment (COLBALT).

Sheet 5: Multiple alignment of cystatin protein sequences using Constraint-based Multiple Alignment (COLBALT).

Sheet 6: Multiple alignment of glutathione s-transferase protein sequences using Constraint-based Multiple Alignment (COLBALT). Multiple alignment of protein sequences using Constraint-based Multiple Alignment (COLBALT).

