## Supplementary figures and images for "Tick Innate Immune Responses to Hematophagy and *Ehrlichia* Infection at Single-Cell Resolution"

### Supplemental Fig. 1.tiff

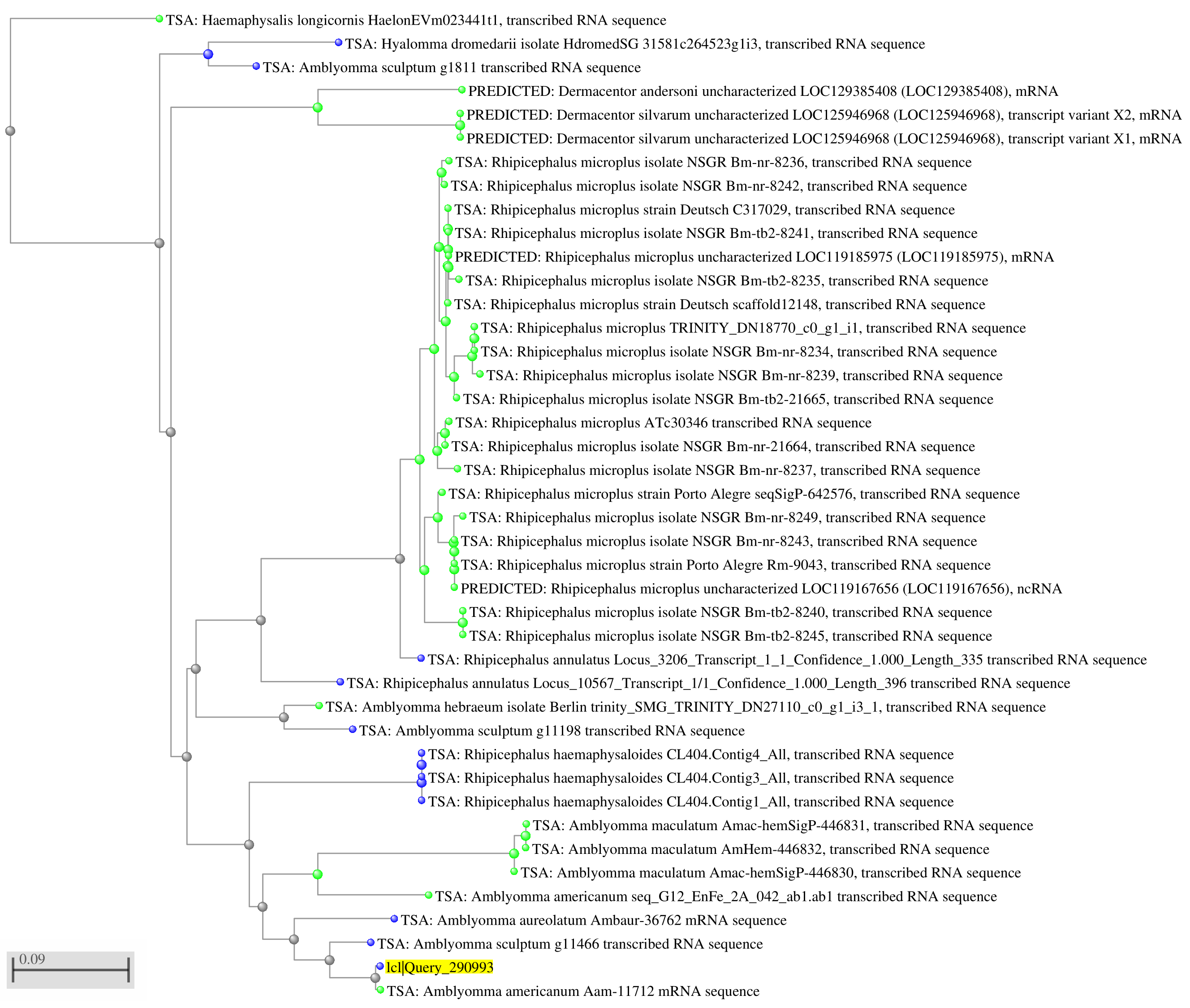

### Supplemental Fig. 2.tiff

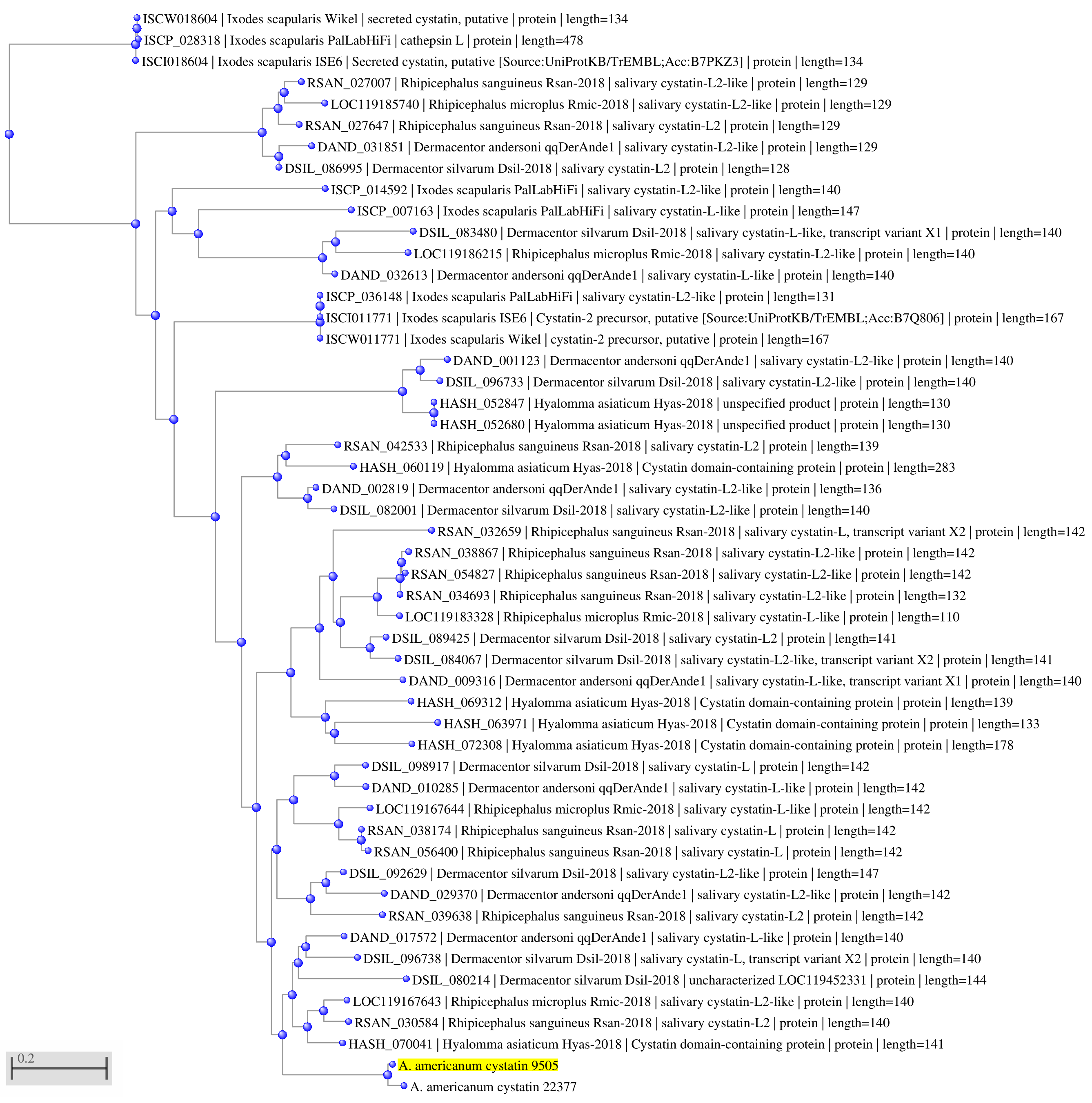

### Supplemental Fig. 3.tiff

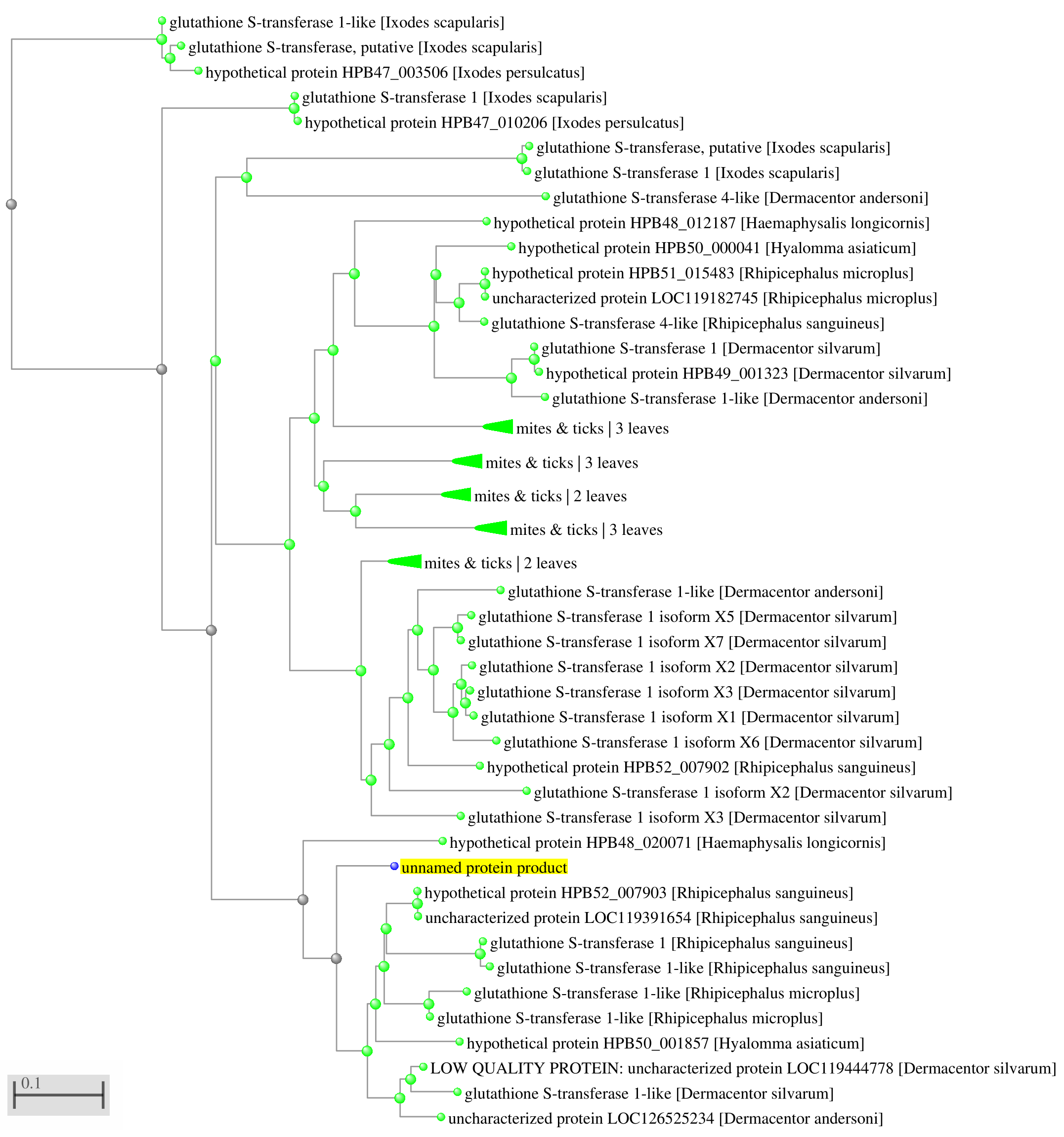
